# Linear indexing for all strings under all internal nodes in suffix trees

**DOI:** 10.1101/2021.10.25.465764

**Authors:** Anas Al-okaily, Abdelghani Tbakhi

## Abstract

Suffix trees are fundamental data structure in stringology. In this work, we introduce two algorithms that index all strings/suffixes under all internal nodes in suffix tree in linear time and space. These indexes can contribute in resolving several strings problems such as DNA sequence analysis and approximate pattern matching problems.

## Introduction

Numerous number of string problems are presented in several scientific fields including biological and medical fields. These problems include exact and approximate pattern matching, motif search, lowest common ancestor, and finding tandem repeats. The inputs for these problems could be small documents, databases, DNA data, or corporational big data. In order to resolve string problems more efficiently, several data structures were designed and are used such as mainly suffix trees [1, 2, 3], suffix arrays [4], and FM-index [5].

Building suffix trees, suffix arrays, and FM indexes needs linear time and space. Given the fact that building suffix trees needs more constant factor compared to suffix arrays and FM-indexes, the design of suffix tree structure is more flexible and dynamic. This is attributed to the fact that the structure of suffix trees allows us to notice many systematic redundancies among the strings in the input data when compared to the structure of suffix arrays and FM indexes. One can easily spot that the set of suffixes under an internal node in the suffix tree for some input data can be equal set, subset, or superset of the set of suffixes under many other internal nodes. This observation is not possible using the structure of suffix arrays and FM indexes. As a result and in order to track and simplify the redundancies in the input data, suffix trees would be exclusively the optimal and most efficient choice. Once these redundancies got tracked and simplified in suffix trees, resolving complex string problems will be easier and more efficient than resoling them using suffix arrays, FM indexes, or even suffix trees in their pure form).

In this work, we introduce two algorithms that index all strings/suffixes under all internal nodes in suffix trees in linear time and space.

## Methods

Let *T* be a text of length *n* derived from alphabet of size Σ and *T*_*i*_ is the suffix at position *i* in *T*. Let *ST* is the suffix tree of *T, Depth*(*n*) is the depth of node *n* i.e. sum of lengths of all edges between root node of *ST* and node *n*, and *Suffix index*(*l*) is the suffix index recorded at leaf node *l*.

### Definition 1

Let *S* be the set of leaf nodes *under* an internal node *x* in *ST*, then suffixes under node *x*, denoted as *SU* (*x*), are the suffixes labeled between node *x* and each leaf node in *S* i.e. SU(x) = {*T*_*Depth*(*x*)+*Suffix Index*(*l*)_ ∀ l ∈ *S*.}.

As an example, suffixes *under* node 20, SU(node 20), in Figure 1 are: “CTAAG$”, “G$”, and “TTTAACTAAG$” (which are suffixes with indexes 13, 17, and 7 respectively). Now, note that when node *a* has a suffix-link to node *b*, then all the suffixes under node (*SU* (*a*)) must be a subset or equal to *SU* (*b*). This indicates that if we assign index values to the suffixes in *SU* (*a*), these index values can implicitly be used/applied to index the same suffixes in *SU* (*b*). Therefore, when indexing the suffixes under node *b*, only the set of *SU* (*b*) − *SU* (*a*) suffixes will be needed to be indexed. This process can be applied recursively so that if node *b* has a suffix-link to node *c*, then we will need to index only suffixes *SU* (*b*) − *SU* (*b*) and nothing will be needed for the set of suffixes *SU* (*c*) − *SU* (*a*) as they have already been indexed. Moreover, as each internal node in the structure of *ST*, say node *x*, may have up to *O*(Σ) nodes with suffix-link to node *x* and in order to build the above indexing schema, we should start by indexing all suffixes under each node with suffix-link to node *x* before indexing node *x* itself (which indicates a postorder traversal process). The need to compute this postorder traversal process urged designing and building the following tree structure name as *OSHR* tree.

**Fig. 1.**
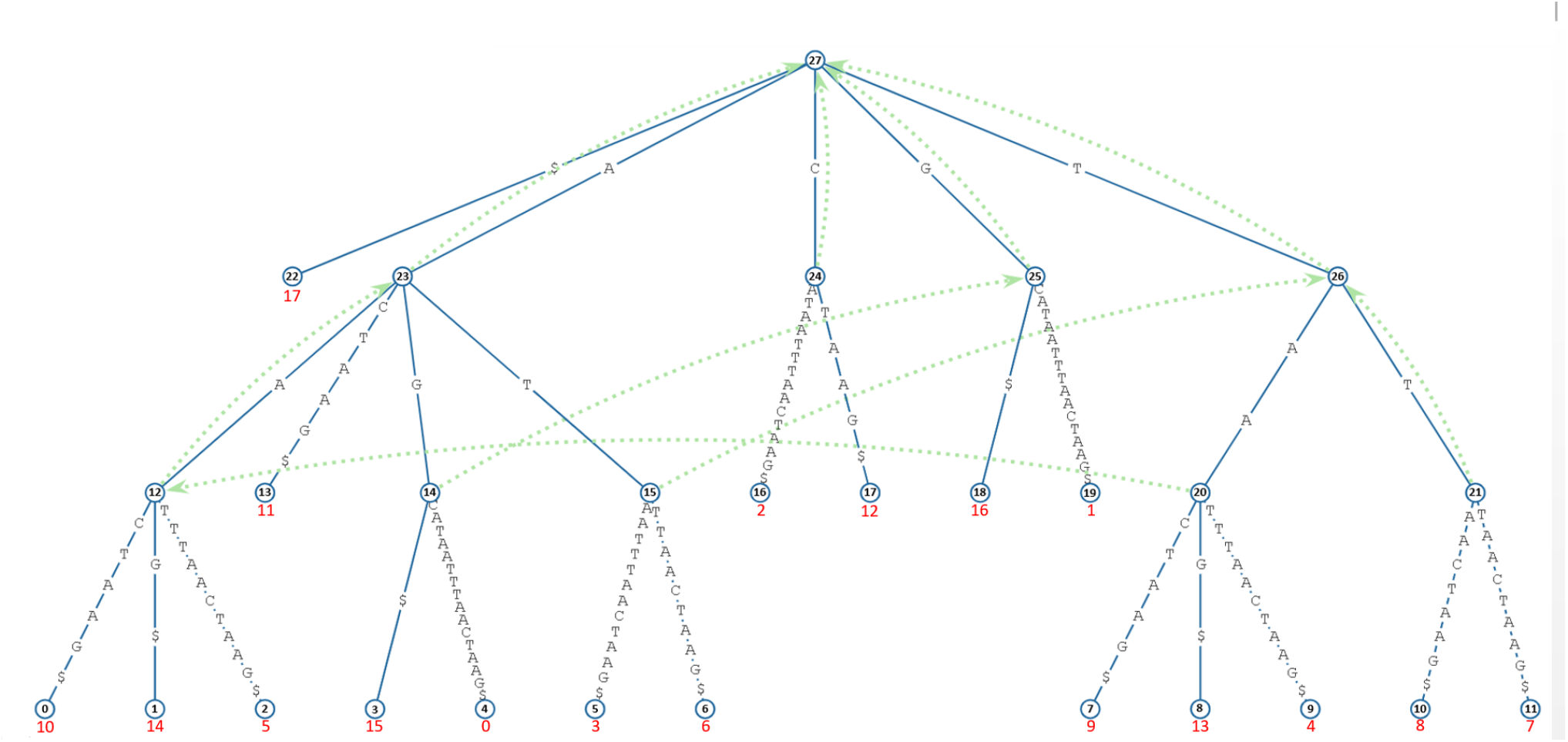
Suffix tree of string AGCCTAATTTAACTAAG$ (drawn at https://hwv.dk/st/?AGCATAATTTAACTAAG$). Suffix indexes start by 1 (not 0)

### *OSHR* tree

Given *ST*, the structure of *OSHR* tree is as the following:

- Root node is the root of *ST*.
- Leaf nodes are all internal nodes in *ST* with no incoming suffix-links. Node 14 from Figure 1 as an example.
- Internal nodes are all the internal nodes in *ST* with at least one incoming suffix-link. Node 25 from Figure 1 as an example.
- Internal nodes are all internal nodes in *ST* with at least one incoming suffix-link (node 25 from Figure 1 as an example).
- Each internal node, let’s say node *x*, must has a list denoted hereafter as *nodes suffix linked to me* to store the nodes suffix-linked to node *x* (the upper bound of length of this list at any internal node is *O*(Σ)). As an example, the list *nodes suffix linked to me* of node 26 in Figure 1 must contain nodes 15 and 21.
- There is a directed edge from node *a* to node *b* if node *b* has a suffix-link to node *a*.
- Edges have no labels.
- Leaf nodes in *ST* are not included in the structure of *OSHR* tree.

The directed edges in *OSHR* tree which are the opposite of suffix-links are actually a simple form of Weiner-links in *ST* (Weiner-link as defined at [6, 7, 8]). Due to the construction properties of *ST* and suffix-links, *OSHR* tree will be a directed acyclic graph. Clearly, the space and time costs for building *OSHR* tree are linear *O*(Σ*n*) and can be constructed implicitly (inside *ST* tree) or explicitly (outside *ST*). The structure of *OSHR* tree is different than suffix-tour-graph [9] and suffix-link-tree [9, 7, 8]. Unlike suffix-tour-graph, *OSHR* structure is acyclic graph. Compared to suffix-links tree structure, the edges in *OSHR* tree have no labels, does not include the leaf nodes of *ST*, and the leaf nodes in *OSHR* tree are internal nodes in *ST* with no incoming suffix links. What the acronym “*OSHR*” stand for is provided in the Acknowledgment section.

### *OT* indexing

In order to track all similarities and redundancies among the strings/suffixes under different internal nodes in *ST*, the following traversal must be performed.

#### Definition 2

Indexing (processing) suffixes/strings under nodes (based on *ST* structure) that have suffix-links to node (*x*) and not re-indexing (re-processing) the same suffixes/strings under node *x* throughout a postorder traversal of *OSHR* tree is referred hereafter as *OT* indexing (processing, the reason for the *OT* acronym is provided in the Acknowledgment section).

A simple example, let node 26 in Figure 1 is the current visited node in postorder traversal of *OSHR* tree, then node 15 and 21 must have been visited already where the following have been already processed at each of them. Suffixes under node 15 which are suffix index 5 (“AATTTAACTAAG$”) and suffix index 8 (“TTAACTAAG$”) must have already been indexed/processed. Under node 21, suffixes with indexes 10 (“AACTAAG$”) and 9 (“TAACTAAG$”) must have been indexed. Now (at node 26), only suffix 16 (“AAG$”) will need to be indexed. This way, all suffixes under node 26 could be indexed without an explicit indexing (processing) all of them. Once postorder traversal of *OSHR* tree reaches the root node, all suffixes under each internal node (including root node) must have been indexed.

#### Algorithm 1

Non-Trivial algorithm for Finding Base Suffixes

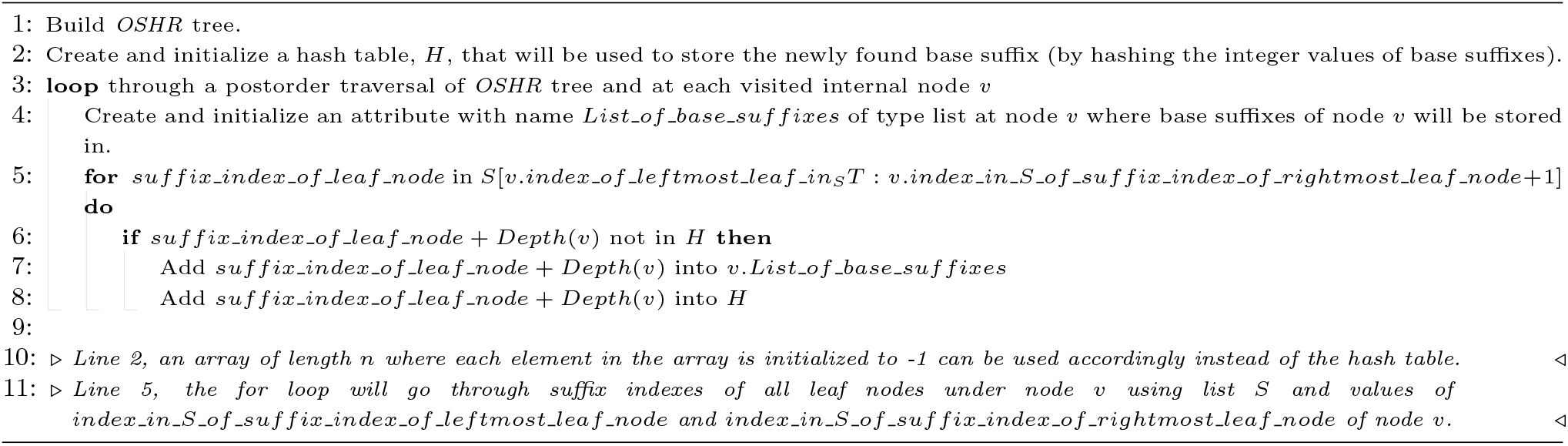

#### Algorithm 2

Non-Trivial algorithm for Finding Base Suffixes

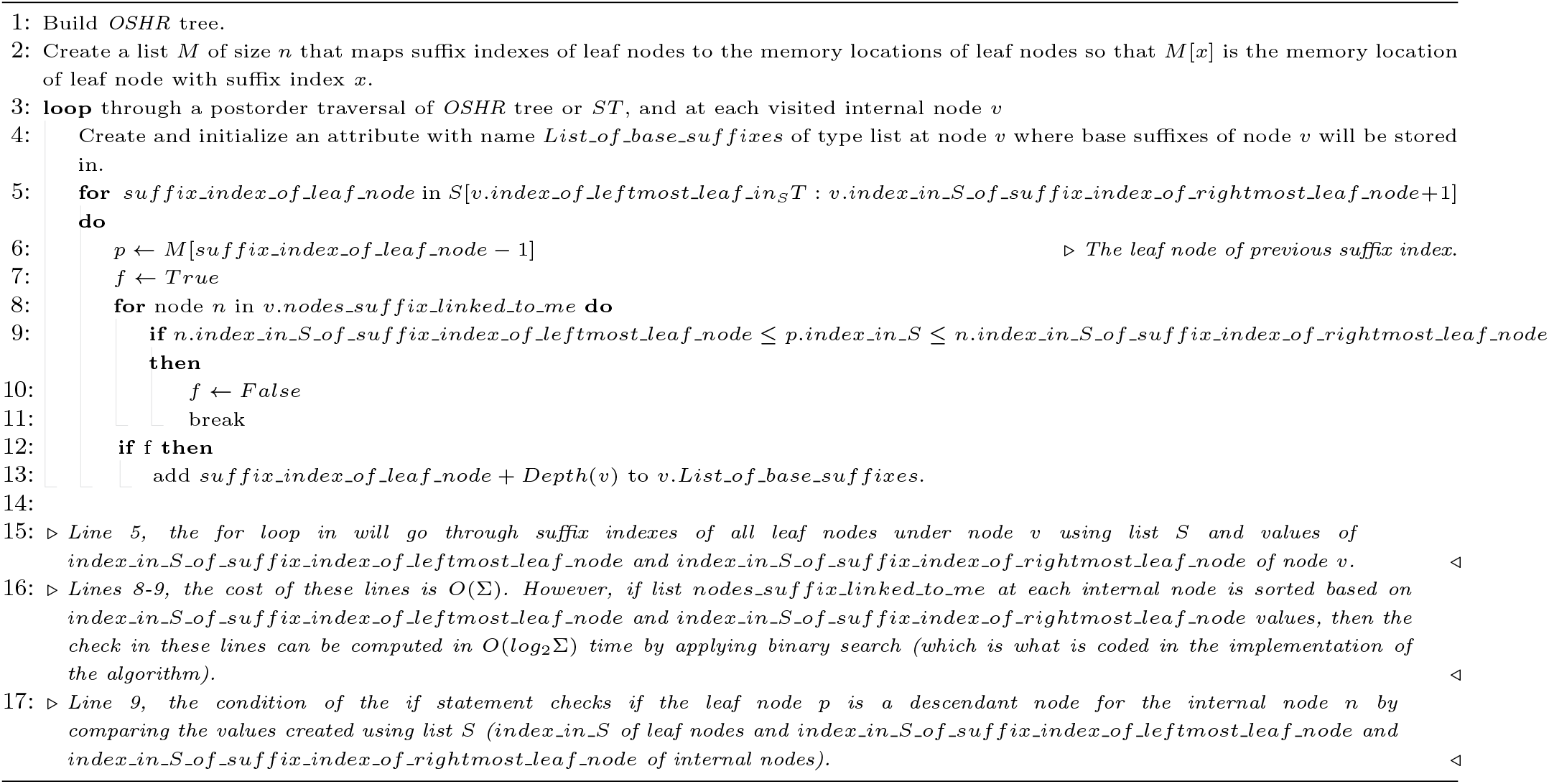

Two types of *OT* indexing will be introduced in this paper, indexing all suffixes under all internal nodes in *ST* in linear costs (*OT* indexing of base suffixes) and indexing all paths between each internal node in *ST* and each of its descendant internal nodes in linear costs (*OT* indexing of base paths).

### Base suffixes

Initially, base suffixes will be defined and later we will describe how to find base suffixes under each internal nodes in *ST* in linear time and space.

#### Definition 3

base suffixes are suffixes that are presented under a node in *ST*, let’s say node *x*, and never have been presented under any of the nodes suffix-linked to node *x*. In case the node is an *OSHR* leaf node, then all suffixes under this node are base suffixes.

The following are examples from Figure 1. The base suffix under node 26 is suffix “AAC$”, the base suffixes for node 23 are “TAATTTAACTAAG$”, “GCATAATTTAACTAAG$”, “ATTTAACTAAG$”, “AG$”, and “ACTAAG$”. As node 20 has no incoming suffix-links, then all suffixes under it are base suffixes. Base suffixes for node 12 are none.

The total number of base suffixes under all internal nodes in *ST* is exactly equal to *n* (as demonstrated and proved in the implementations). The term “base” is selected to indicate that this is the first appearance of this suffix under any internal node.

#### Definition 4

If *s* is a base suffix under node *x*, then *extent suffixes* of base suffix *s* are each suffix under each node that is ancestor to node *x* based on *OSHR* tree structure (throughout suffix-links) where each of these suffixes is equal to suffix *s*.

For example, the suffix “TTAACTAAG$” which ends at leaf node 6 is a base suffix under node 15, the extant suffixes for this base suffix are: suffix “TTAACTAAG$” under node 26 which ends at leaf node 11, and the one under root node which ends at leaf node 9.

The upper bound for the number of extent suffixes for any base suffix is *O*(*h*) where *h* is the height of the *ST*. The last extent suffix of any base suffix is the one under the root node.

#### Algorithm 3

Linear algorithm for Finding Base Suffixes

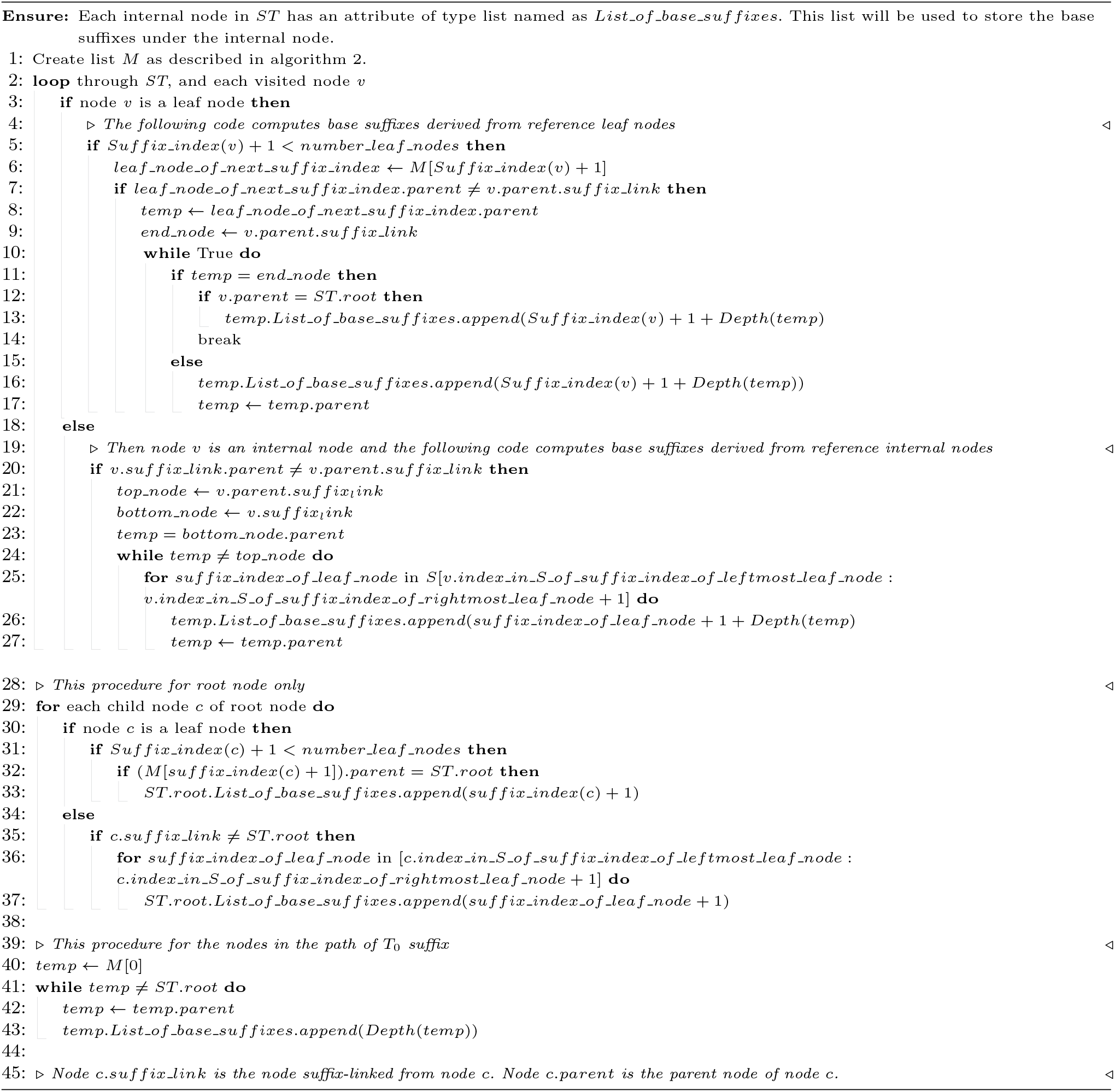

This way, once a base suffix is processed/indexed, this processing/indexing can be applied *implicitly* to all *O*(*h*) extent suffixes. Hence, *OT* indexing/processing all suffixes under all internal nodes in *ST* can be computed with a factor of *n*.

### Finding base suffixes

Base suffixes under each internal nodes can be computed throughout a postorder traversal of *OSHR* tree or *ST* tree where at each visited internal node, *v*, check whether each suffix under node *v* presented or not under any of the Σ nodes suffix-linked to node *v*. If not, then this suffix is a base suffix for node *v*. The cost of this process will be *O*(*n*) space (for storing the base suffixes at each internal node) and *O*(*nhc*) time where *nh* is the sum of looping through all suffixes under all internal nodes in *ST* and *c* is the time cost for the checking procedure.

Trivially, the checking can be performed by looping through each suffix under each of the Σ nodes suffix-linked to node *v*, however this will be very costly (*O*(*n*^2^*h*)). We will now describe non-trivial algorithms with *O*(*nh*), non-trivial algorithms with *O*(*nhlog*_2_Σ) time costs, and then describe a linear algorithm.

Before of all, *ST* must be already built. In addition, the following list named as *S* must be built which will be used in the three algorithms. Traverse *ST* and append the suffix indexes of leaf nodes from left to right into list *S* (the first element of *S* is the suffix index of leftmost leaf node and the last element is the suffix index of rightmost leaf node). Meanwhile, create an attribute named as *index in S* at leaf node where its value is the index of leaf node in list *S*, and create recursively two attributes at each internal node: *index in S of suffix index of leftmost leaf node* and *index in S of suffix index of leftmost leaf node* where the value of these attributes are the index in *S* of leftmost and rightmost leaf nodes under the internal node, respectively.

#### Algorithm 4

Non-Trivial algorithm for Finding Base Paths

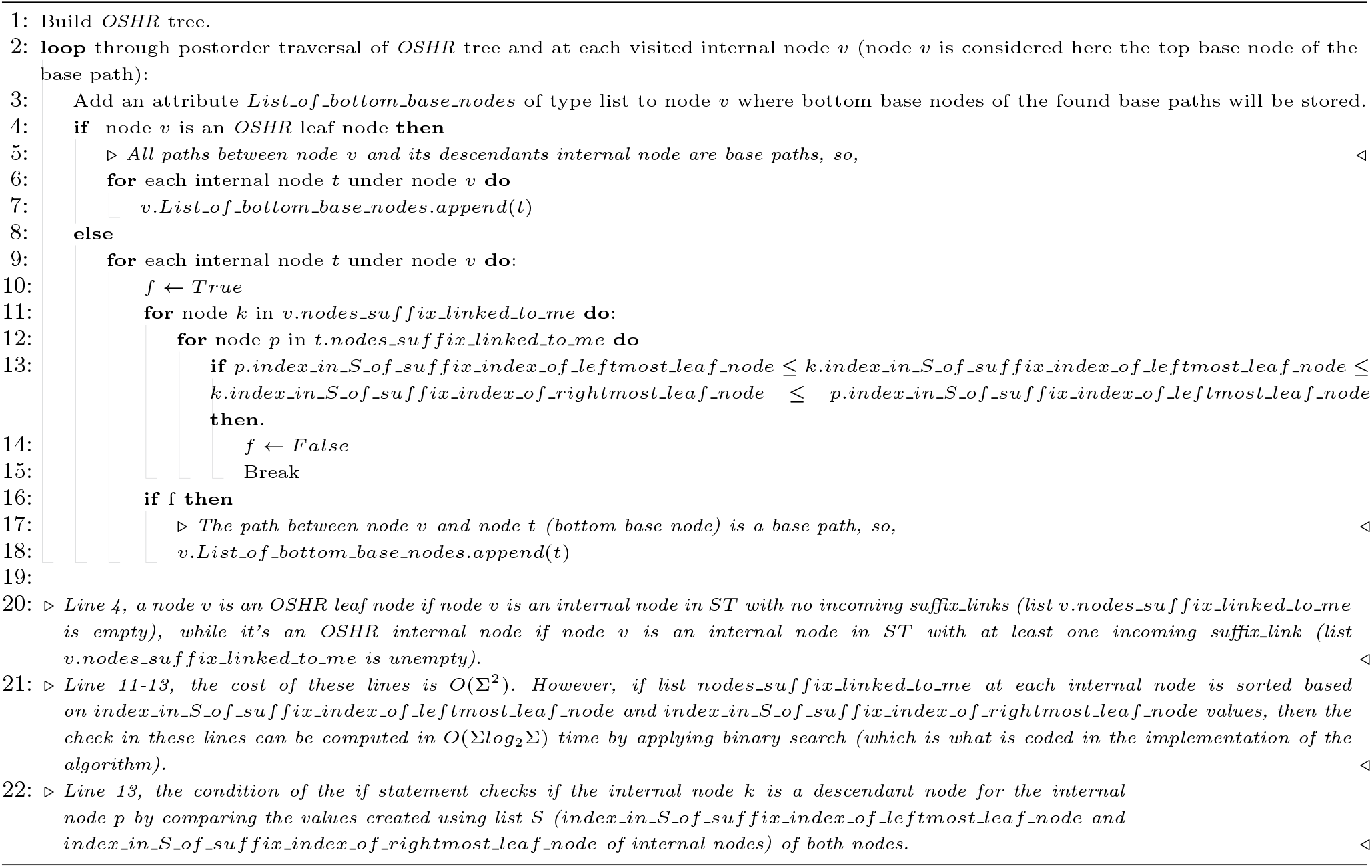

The first non-trivial algorithm will cost *O*(*nh*) with auxiliary *O*(*n*) space (hash table) as shown in Algorithm 1. The second non-trivial algorithm will cost *O*(*nhlog*_2_Σ) with auxiliary *O*(*n*) space as shown in Algorithm 2.

The linear algorithm motivated from the fact that the total number of base suffixes for all internal nodes in *ST* is *n*, then finding each base suffix in constant time will cost *O*(*n*) time. Before stating this algorithm, we need to define the followings.

#### Definition 5

Let A be a leaf node in *ST* with suffix index *x*, B is the parent node of node A, C is the leaf node with suffix index *x* + 1, and D is the parent node of node C. If the node suffix-linked from node B is not node D, then denote node A as a *reference leaf node* and denote each node between node C and node D as an *inbetween* node.

As an example from Figure 1, node 6 is reference leaf node for node 21 and node 21 is an inbetween node for node 6. Note that a reference leaf node can have more than one inbetween node and an inbetween node can have up to *O*(Σ) reference leaf nodes. In addition, the total number of reference leaf nodes for all internal nodes in *ST* cannot be more than *n*.

#### Definition 6

Let A be an internal node in *ST*, B is the parent node of node A, C is the suffix-link node of node A, and D is the parent node of node C. If the node suffix-linked from node B is not node D, then denote node A as a *reference internal node* and denote each node between node C and node D as an *inbetween* node.

As an example from Figure 1, node 20 is reference internal node and node 23 is an inbetween node for node 20. Note that a reference internal node can have more than one inbetween node and an inbetween node can have up to *O*(Σ) reference internal nodes. In addition, the total number of reference internal nodes for all internal nodes in *ST* is much less than *n*. Moreover, an inbetween node can be derived from reference leaf node (or more) and reference internal node (or more).

Now, the linear algorithm can be computed with or without explicit finding and recording of inbetween nodes and their reference internal nodes and/or reference leaf nodes. The main form of this algorithm is without explicit finding and recording of inbetween nodes. Algorithm 3 states this algorithm. The algorithm that explicitly create and record inbetween nodes and their reference leaf nodes and reference internal nodes is provided in the code provided in this work.

As the upper bound of the number of reference leaf nodes and the upper bound of the number of reference internal nodes is Σ for any internal node (given that most of internal nodes will not be inbetween nodes in the first place), the cost for finding inbetween nodes will be *O*(Σ*n*). For the computing base suffixes, the cost will be *O*(*n*) as the total number of base suffixes under all internal nodes in *ST* is equal to *n*.

#### Theorem 1

Finding all base suffixes under all internal nodes in *ST* can be computed in linear time and space (*O*(*n*)).

Consequently, once base suffixes are computed and found under each internal nodes in *ST* and throughout *OT* indexing/processing, any index/process *P* with cost *p* for a base suffix *s* under an internal node will implicitly be applied to all *O*(*h*) extent suffixes of *s*. Hence, the total cost of applying process *P* for all suffixes under all internal nodes in *ST* will be to the factor of *n* and will cost *O*(*pn*) instead of (*phn*).

### Base paths

The motivations behind this index are the following observations. Firstly, the main complexity in a tree structure is due to the branching caused by internal nodes. Secondly, the tails of suffixes (label between leaf node and its parent node) are usually of very long length, hence processing them are costly. Thirdly, assuming a process reaches an internal node where all its children are leaf nodes, then the cost for performing some computations on these leaf nodes will be bounded to Σ factor. Therefore, there is no need to always perform any indexing/processing for the tails of the suffixes and it will be sufficiently efficient for many applications to perform indexing/processing only to the labels/paths between each internal node and its descendant *internal* nodes.

#### Definition 7

A path between two internal nodes, let’s say nodes *a* and *b*, where this path was never appeared between any other two internal nodes throughout suffix-links to node *a* and to node *b* are referred hereafter as *base path*. This is induced by having no incoming suffix-link to node *a, b*, both, or there are incoming suffix-links to node *a* and to node *b* but the label between any node with suffix-link to node *a* and any node with suffix-link to node *b* is not equal to the label between node *a* and node *b*. Hereafter, node *a* is referred as *top base node* and node *b* as *bottom base node* for the base path.

For example, the path in Figure 1 (given that the input text for the *ST* in Figure 1 is very small) between node 23 and node 14 is a base path as it was never appeared between any other two internal nodes. Paths between nodes 26 and 20 and path between nodes 23 and 12 are also base paths.

#### Definition 8

If *p* is base path between top base node *a* and bottom base node *b*, then the path between node *a*^′^ (node suffix-linked from node *a*) and node *b*^′^ (node suffix-linked from node *b*) is an *extent path* for the base path *p*. This applies between all ancestor nodes (under *OSHR* tree structure and throughout suffix-links) of node *a* and node *b*. The last extent path for base path *p*, is the one where the top base node is the root node of *ST*.

As an example, the path between the root node and node 25 is an extent path for the base path between nodes 23 and 14.

Note that the set of unique labels of all base paths *must* eventually equal exactly to the set of labels of paths between the root node and all internal nodes in *ST*. This must be true, as proved also by implementation, since a path between the root node and a descendant internal nodes is the last appearance (extent) path of some base path/s throughout suffix-links in *ST*. As any path between the root node and internal node can have up to *O*(Σ) paths from other pair of nodes, the total number of base paths can’t be more than *O*(Σ*n*).

### Finding base paths

All base paths in *ST* can be found using a trivial algorithm with time cost of *O*(*hn*^2^) as follows. Perform postorder traversal of OSHR tree and at each visited node *v*: check if the label of each path between node *v* and a descendant internal node of *v* is presented between any node (with suffix-link to node *v*) and one of its descendant internal node; if not, then this path is a base path. However, in this work we will introduce a non-trivial algorithm that costs *O*(*nh*Σ*log*_2_Σ) time and a linear algorithm (*O*(Σ*n*) time and space).

The first non-trivial algorithm for finding base suffixes which uses hash table can’t be applied here to find base paths as base paths are not totally unique like base suffixes (total number of base suffixes are exactly *n* where for base paths they can be up to Σ*n*). For instance, two sibling nodes under *OSHR* tree structure (both nodes must suffix-link to the same node), can have base path under each of them with the same label. So if the base path under the first node is found and hashed, then the base path under the second node will not be detected and found as it was found/hashed already. However, Algorithm 4 states non-trivial algorithm which is similar to the second non-trivial algorithm for finding base suffixes can compute base paths with *O*(*nh*Σ*log*_2_Σ) time and *O*(Σ*n*) space.

As the total number of base paths is no more than Σ*n*, if the time cost for finding each base path is constant, then finding all base paths will cost *O*(Σ*n*) time and space. Certainly this is achievable using Algorithm 5.

#### Theorem 2

Finding all base paths under each internal nodes in *ST* can be computed in linear time and space (*O*(Σ*n*)). Hereby, once base paths are computed and found under each internal node in *ST*, throughout *OT* processing any processing *P* with cost *p* for a base path *t* under an internal node will implicitly be applied to the *O*(*h*) extent paths of *t* under the other internal nodes. Hence, the total cost of applying process *P* for all paths under all internal nodes in *ST* will be to the factor of *n* and will cost *O*(*np*) instead of (*phn*).

## Results

In order to test the proposed algorithms, we used the genomes: WS1 bacterium JGI 0000059-K21 (Bacteria), Astrammina rara (Protist), Nosema ceranae (Fungus), Cryptosporidium parvumIowa II (Protist), Spironucleus salmonicida (Protist), Tieghemostelium lacteum (Protist), Fusarium graminearumPH-1 (Fungus), Salpingoeca rosetta (Protist), and Chondrus crispus (Algae).

As a prepossessing step, header lines and newlines were removed from the fasta file and any small letter nucleotide were capitalized. This generates a one line genome file for each fasta file where all nucleotide are capital. The python script used for this preprocessing step is provided at https://github.com/aalokaily/Finding_base_suffixes_and_base_paths.

All five algorithms stated in this work were implemented using Python language and are available at https://github.com/aalokaily/Finding_base_suffixes_and_base_paths. For base suffixes finding, the outcomes of the non-trivial algorithm (algorithm 1) and linear algorithm (algorithm 3) were reported in order to validate the finding of base suffixes and compare their time costs. As the non-trivial algorithm (algorithm 2) is theoretically and as was found practically (data not shown) slower than algorithm 1, we ignore its comparisons. For base paths finding, the outcomes of non-trivial algorithm (algorithm 4) and linear algorithm (algorithm 5) were reported in order to validate the finding of base paths and compare their time costs. All results are shown in Table 1.

**Table 1.** Results for testing, validating, and comparing algorithms 1 and 3 for finding base suffixes and algorithms 4 and 5 for finding base paths.

| Genome | WS1 bacterium | Astrammina | Nosema | Cryptosporidium | Spironucleus | Tieghemostellium | Fusarium | Salpingoeca | Chondrus |
| --- | --- | --- | --- | --- | --- | --- | --- | --- | --- |
| GenBank Accession | GCA000398605.1 | GCA000211355.2 | GCA000988165.1 | GCA000165345.1 | GCA000497125.1 | GCA001606155.1 | GCF000240135.3 | GCA000188695.1 | GCA000350225.2 |
| No. of alphabets | 5 | 5 | 4 | 15 | 5 | 4 | 5 | 5 | 5 |
| No. of nuc/leaf nodes | 509,552 | 1,450,096 | 5,690,749 | 9,102,325 | 12,954,589 | 23,375,663 | 36,458,047 | 55,440,310 | 104,980,421 |
| No. of Internal nodes | 328,917 | 926,087 | 3,849,880 | 5,931,181 | 8,684,514 | 15,606,221 | 22,972,065 | 37,480,799 | 81,909,252 |
| No. of OSHR leaf nodes | 132,774 | 380,518 | 1,353,289 | 2,328,480 | 3,127,031 | 5,740,807 | 9,800,385 | 11,545,032 | 15,307,982 |
| No. of OSHR internal nodes | 196,143 | 545,569 | 2,496,591 | 3,602,701 | 5,557,483 | 9,865,414 | 13,171,680 | 25,935,767 | 66,601,270 |
| Alg 1 (time complexity/sec) | 2,409,867/3 | 6,838,684/9 | 26,549,891/37 | 42,989,315/61 | 60,212,775/92 | 109,641,901/172 | 172,751,172/278 | 250,969,523/446 | 457,348,511/983 |
| Alg 3 (time complexity/sec) | 6,117,025/3 | 18,036,912/8 | 83,860,995/41 | 186,083,328/113 | 215,252,439/110 | 364,737,747/209 | 851,871,768/561 | 19,667,387,766/8,763 | 2,787,481,705/1,508 |
| No. of base paths | 1,229,296 | 3,685,339 | 16,277,962 | 27,440,361 | 37,673,155 | 73,606,066 | 116,915,929 | 161,201,632 | 234,284,261 |
| Alg 4 (time complexity/sec) | 3,940,965/6 | 11,336,761/19 | 47,652,774/87 | 76,241,139/140 | 108,625,373/214 | 201,220,732/396 | 307,913,746/647 | 479,928,226/1,070 | 882,404,297/3,222 |
| Alg 5 (time complexity/sec) | 6,314,996/21 | 18,734,271/63 | 99,497,879/328 | 210,245,283/829 | 238,310,327/877 | 430,212,888/1,542 | 804,854,850/3,766 | 15,946,058,214/93,295 | 3,670,449,444/14,452 |

First of all, all base suffixes and base paths found under each internal nodes in *ST* using the linear algorithms (algorithms 3 and 5) were the same as the ones found using their correspondent non-trivial algorithms (algorithm 1 and 4). Clearly as shown in Table 1, both linear algorithms are faster than the correspondent non-trivial algorithms. In addition, the cost for linear algorithms from small size to large size genomes grows in constant factor as expected while their respected non-trivial algorithms grows larger as they are dependent on a variable factor (the *h* value). Lastly, the total number of base paths for any genome (small to large) is no more than *O*(Σ*n*).

### Algorithm 5

Linear algorithm for Finding Base Paths

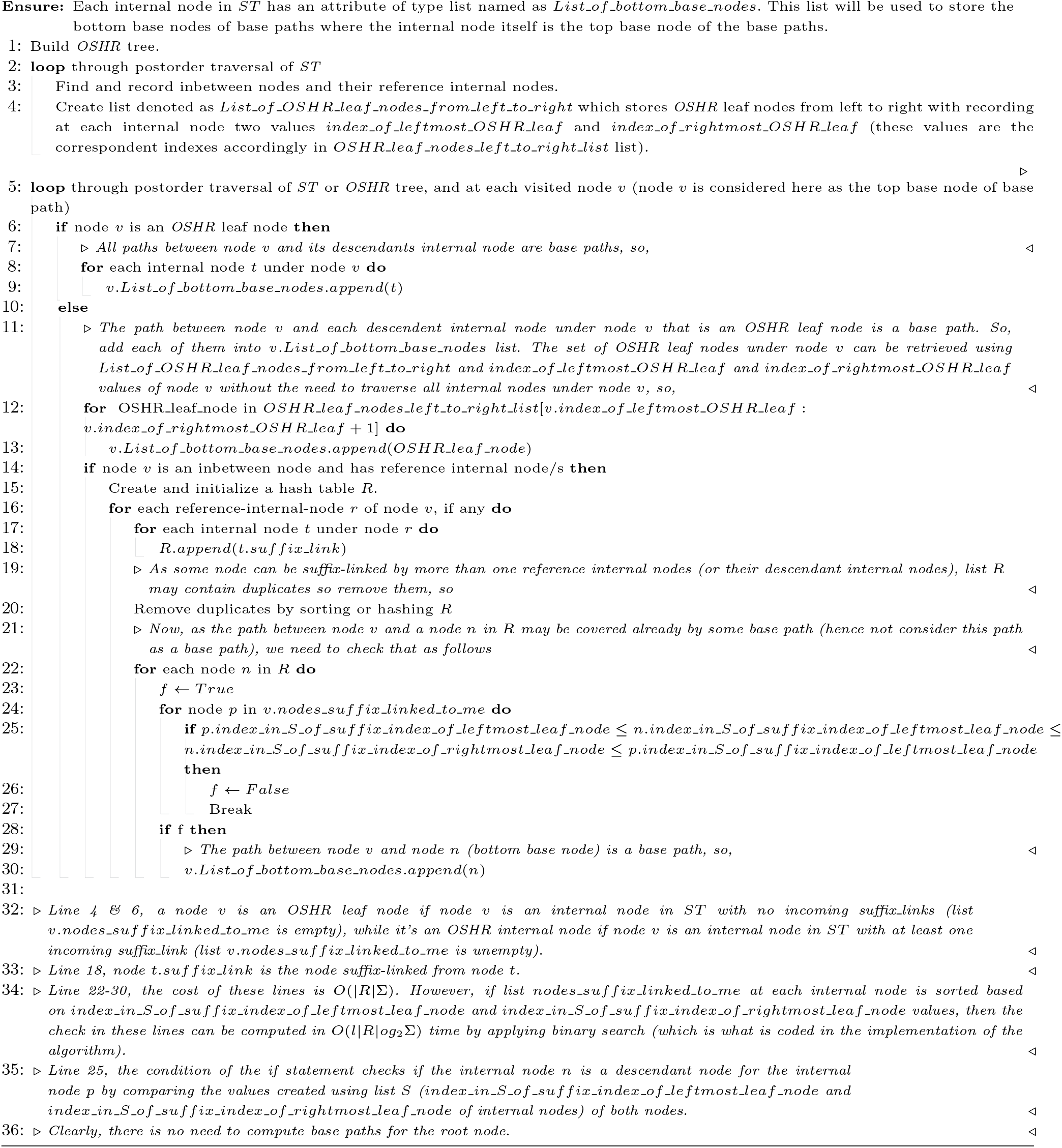

## Acknowledgments

The name “OSHR” tree stands for Okaily-Sheehy-Huang-Rajasekaran. These letters stand for the last name of the first author and last names of his PhD committee in the University of Connecticut, Department of Computer Science, as the initial version of the error tree structure was a chapter in the PhD dissertation of the first author. The committee members were: Chun-Hsi Huang (Major Advisor), Sanguthevar Rajasekaran, and Don Sheehy. This naming meant to tribute and appreciate their kind, influential, and professional teaching and supervision. The name OT stands for the last names of the authors of this work.

## Data and materials availability

The algorithms proposed in this paper are implemented using python programming language and are available at https://github.com/aalokaily/Finding_base_suffixes_and_base_paths.

